# Diverse patterns of intra-host genetic diversity in chronically infected SARS-CoV-2 patients

**DOI:** 10.1101/2024.11.23.624482

**Authors:** Natalie Rutsinsky, Adi Ben Zvi, Ido Fabian, Shir T. Segev, Bar Jacobi, Sheri Harari, Suzy Meijer, Yael Paran, Adi Stern

## Abstract

In rare individuals with a severely immunocompromised system, chronic infections of SARS-CoV-2 may develop, where the virus replicates in the body for months. Sequencing of some chronic infections has uncovered dramatic adaptive evolution and fixation of mutations reminiscent of lineage-defining mutations of variants of concern (VOCs). This has led to the prevailing hypothesis that VOCs emerged from chronic infections. To examine the mutation dynamics and intra-host genomic diversity of SARS-CoV-2 during chronic infections, we focused on a cohort of nine immunocompromised individuals with chronic infections and performed longitudinal sequencing of viral genomes. We show that sequencing errors may cause erroneous inference of high genetic diversity, and to overcome this we used duplicate sequencing across patients and time-points, allowing us to distinguish errors from low frequency mutations. We further find recurrent low frequency mutations that we flag as most likely sequencing errors. This stringent approach allowed us to reliably infer low frequency mutations and their dynamics across time. We inferred a synonymous divergence rate of the virus of ∼2×10^−6^ mutations/base/day, consistent with the SARS-CoV-2 mutation rate estimated in tissue culture. The rate of non-synonymous divergence varied widely among the different patients. We highlight two patients with opposing patterns: in one patient the rate of divergence was zero, yet this patient harbored multiple presumably defective viruses at low frequencies throughout the infection. Another patient exhibited dramatic adaptive evolution, including clonal competition. Overall, our results suggest that the emergence of highly divergent variants from chronic infections is likely a very rare event and this emphasizes the need to better understand the conditions that allow such emergence events.

## Introduction

The evolution of SARS-CoV-2 has been marked by the periodic emergence of variants that dramatically differ genetically from the variants circulating at the time. Some of these variants have shown increased transmissibility compared to earlier ones, resulting in rapid replacement of one variant with another (Markov, et al. 2023). These variants were also denoted as Variants of Concern (VOCs), identified by the world health organization due to their heightened risk to public health. One of the leading hypotheses regarding the origin of highly divergent SARS-CoV-2 variants is that they arise in chronically infected individuals (Rambaut and al. 2020; Dennehy, et al. 2022; Harari, et al. 2022; Tay, et al. 2022). Chronic infections are defined as cases where actively replicating virus persists for more than 21 days, and de facto such infections often last months to over a year (Avanzato, et al. 2020; Baang, et al. 2021; Kemp, et al. 2021; Harari, et al. 2022). This is opposed to acute infections, that are normally resolved with a few days to weeks. Chronic infections have been most often observed in immunocompromised individuals, specifically those with hematologic cancer, AIDS, transplant recipients, or autoimmune disorders (Fung and Babik 2021; DeWolf, et al. 2022).

The possible role chronic SARS-CoV-2 infections may have in generating VOCs has led to a heightened research interest in these patients. Such chronic infections are thought to be rare, as the proportion of immunocompromised individuals in the general population is low (Antinori and Bausch-Jurken 2023). Notably, a recent study has shown that persistent infections with rebounding viral loads may also be present (but rare) in the general population as well, in individuals not necessarily immunocompromised (Ghafari, Hall, et al. 2024). Having said that, we and others have shown that most chronic infections do not likely lead to onwards transmission (Harari, et al. 2022; Harari, et al. 2024). However, even very rare events, i.e., the emergence of a transmissible variant from a chronically infected patient, may pan out when case numbers were or are high globally.

While many case reports have focused on very rapid evolution occurring in chronic infections (e.g., Choi, et al. 2020; Kemp, et al. 2021), it is not clear if this is the rule or the exception. Moreover, while rapid evolution in these patients may result from strong selection pressures, it could also stem from an increased viral mutation rate specific to these patients. These enigmas require an in-depth investigation of within-host genetic diversity that accumulated during chronic infections. However, one of the major challenges in assessing such within-host genetic diversity are high error rates associated with sequencing low biomass samples of viruses (McCrone and Lauring 2016; Zhao and Illingworth 2019; Gelbart, et al. 2020; Tonkin-Hill, et al. 2021; Valesano, et al. 2021). This makes inferences of levels of genetic diversity, and measurement of mutation frequencies (also denoted as intra-host single-nucleotides variants, iSNVs), very challenging. This has led to several commonly used practices, such as filtering for high frequency mutations (Harari, et al. 2022), filtering for high coverage (Zhang, et al. 2022), or sequencing samples in duplicates (e.g., Tonkin-Hill, et al. 2021; Bendall, et al. 2023). Of note, filtering strategies will tend to lose information, with a tendency to remove, often by definition, low frequency mutations.

We focused here on obtaining reliable sequencing from longitudinal samples of nine chronically infected patients, with the aim of minimal filtering while retaining a biological signal. To this end, we performed duplicate sequencing of each sample and showed that this considerably improves inferences of within-host genetic diversity. We went on to infer patterns of mutation frequencies across time, revealing large differences among patients. This allowed us to infer the synonymous and non-synonymous divergence rates. We show that the synonymous divergence rate across all patients matches previous estimates of the viral mutation rate and substitution rate, obtained from tissue culture experiments and from globally available data, respectively.

On the other hand, the non-synonymous divergence rates varied widely across patients. We highlight the two extreme examples observed in our cohort and go on to discuss the implications of this work to the risk of chronic infections generating new transmissible variants.

## Results

We focused on a set of nine patients of chronically infected individuals who were hospitalized in Tel Aviv Sourasky Medical Center (TASMC), for which we had longitudinal samples available, spanning from 21 to 241 days of infections, and for which we were able to perform independent deep sequencing of almost all samples twice (Methods). Seven of these patients were previously described (Harari, et al. 2022; Meijer, et al. 2024) but were sequenced in depth and in duplicate for the first time herein (Table 1, Methods). In parallel, we compared our results to a set of acutely infected patients previously described (Bendall, et al. 2023).

**Table 1.** Summary of all 9 patients with chronic SARS-CoV-2 infections.

| Patient ID | Background condition | COVID-19 initial detection | Sampling start | Days of sequenced infection | Sex | Age range | ID in previous publication | Pango lineage |
| --- | --- | --- | --- | --- | --- | --- | --- | --- |
| **N1** | Chronic lymphocytic leukaemia (CLL) | 17/02/2022 | 07/03/2022 | 241 | male | 70-79 | P7 (Meijer, et al. 2024) | BA.2 |
| **N2** | Multiple myeloma (MM) | 15/02/2022 | 16/03/2022 | 105 | male | 70-79 | P8 (Meijer, et al. 2024) | BA.2.31 |
| **N3** | Non-Hodgkin lymphoma (NHL) | 07/02/2022 | 11/04/2022 | 76 | female | 70-79 | P10 (Meijer, et al. 2024) | BA.1 |
| **N4** | NHL | 14/04/2022 | 14/04/2022 | 27 | female | 70-79 | P13 (Meijer, et al. 2024) | BA.2 |
| **N7** | CLL | 17/02/2022 | 30/03/2022 | 21 | male | 60-69 | none | BA.1.1 |
| **N8** | CLL | 28/03/2022 | 01/04/2022 | 37 | female | 50-59 | none | BA.1 |
| **P3** | Hodgkin's lymphoma | 01/01/2021 | 01/02/2021 | 80 | male | 30-39 | P3 (Harari, et al. 2022) | B.1.362 (non-VOC) |
| **P4** | Autoimmune skin disease | 01/01/2021 | 27/01/2021 | 35 | female | 70-79 | P4 (Harari, et al. 2022) | B.1.1.7 |
| **P5** | Adult acute lymphoblastic leukaemia (ALL) | 01/10/2020 | 06/10/2020 | 68 | male | 0-9 | P5 (Harari, et al. 2022) | B.1.1.50 (non-VOC) |

We began by testing for proportions of mutations at different codon positions given different frequency thresholds, using single (non-replicated) sequencing of samples. Our results showed equal proportions of mutations at positions 1, 2, and 3 of codons under low frequency thresholds, consistent with sequencing errors and lack of biological signal. Mutations with frequencies higher than 0.01 showed higher prevalence at codon position 3, in line with lower selection at this position, and suggesting mutations called above this threshold are biologically relevant (Fig. S1).However, even following filtering for *f* > 0.01, we show here that there still remains a spurious positive correlation between Ct values and the inferred nucleotide (*π*) diversity in each sample (Methods), and this was true for both acute and chronic infection samples (Fig. S2). In other words, we observed higher diversity in low viral load samples, and there is no reasonable biological justification for such an observation. In line with previous work, we conclude it is much more likely that low viral load samples, where template counts are low, may harbor an error generated during early stages of library preparation, that is then carried on to high frequencies. This spurious correlation was most notable at Ct values higher than 24. Unfortunately, most samples we obtained herein (31 of 42) were at Ct values higher than 24 (Table S1), leading us to perform duplicate sequencing. We developed a simple pipeline that allows inference of mutation frequencies that agree in the duplicates (Methods, Fig. S3).

Following this filtering procedure, the spurious correlation observed previously, reassuringly disappeared (Fig. 1, green line). Indeed, we observed the exact same pattern when analyzing the acute infection data, where accounting for replicates led to loss of the spurious correlation with Ct (Fig. S2). We went on to compare *π* diversity between acute infection and chronic infection samples. Our results showed that on average, *π* diversity is significantly higher in the chronic infections (t-test, p<0.001), consistent with the latter infections offering more time for diversity to accumulate and possibly a higher mutation rate and/or more opportunities for positive selection to increase mutation frequencies, as we go on to explore below.

**Figure 1.**
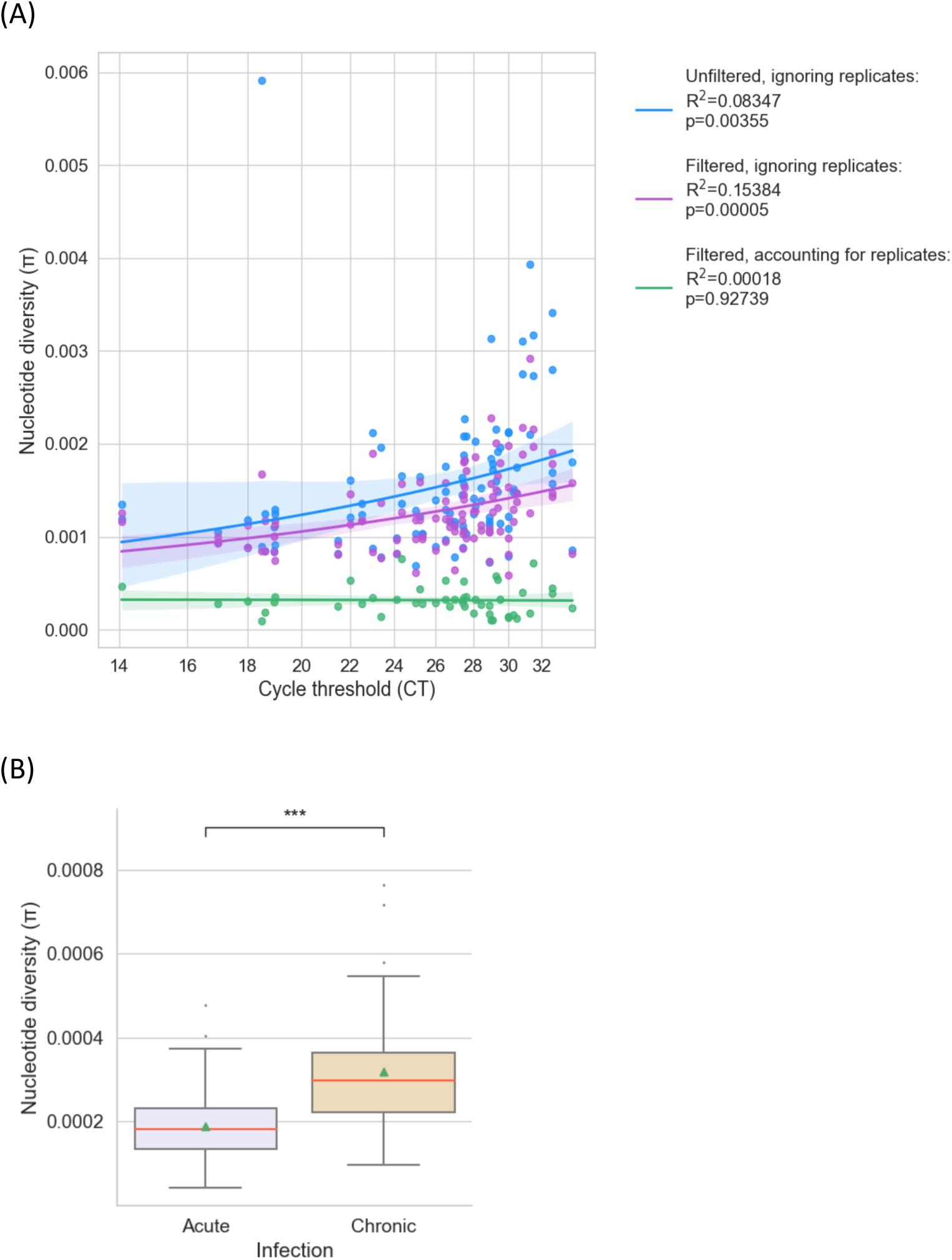
Nucleotide diversity in SARS-CoV-2 samples. (A) Shown is the relationship between Ct, an inverse measure of viral load, and pi diversity of samples from chronic infections. A spurious significant association is obtained when the data is unfiltered and ignoring technical replicates (R^2^=0.08, P<0.005), and is also obtained when filtering of mutations is performed for *f*>0.01, mutation count>50, coverage> 100 (R^2^=0.15, P<0.0001). This association disappears when mutations are called based on their presence in replicate sequencing data (R^2^=0.0, P=0.9). (B) After accounting for replicates, significantly higher diversity is observed in chronic versus acute infection samples (p<0.001, t-test).

We next set out to infer patterns of mutation frequencies across time. In our initial analysis, we observed some mutations present in many patients at a constant and low frequency (*f*=∼1-10%) across time (Fig. S4). This constant pattern was suspicious, as it occurred concurrently with some mutations increasing dramatically in frequency and fixing. We suspected that these mutations may reflect consistent sequencing errors associated with the ARTIC sequencing method we used, which was very widely used during the pandemic. Notably, our analysis filters out substitutions that were flagged at early stages of the pandemic as “suspicious positions” (Methods). However, this list was based on substitutions (i.e., *f*=∼100%), whereas our focus here is on minor variants. To this end, we analyzed a set of acute infections (Bendall, et al. 2023) and generated a set of “suspicious minor mutations” that were present across 20% or more of samples. Such mutations could be driven by selection that occurs in samples in parallel, or due to consistent sequencing errors. We performed a thorough analysis of this set, in which we did not detect a biological signal but rather that this set is likely enriched for sequencing errors. This conclusion was driven by lack of enrichment in spike, lack of enrichment for synonymous or non-synonymous mutations, yet adjacency to primers (Fig. S5, S6; Methods). Although we cannot rule out that the parallel occurrence of some of these mutations may be due to selection, these are likely a minority. All mutations in this set were filtered from our results, allowing us to focus on reliable mutations and their patterns across time (Fig. 2).

**Figure 2.**
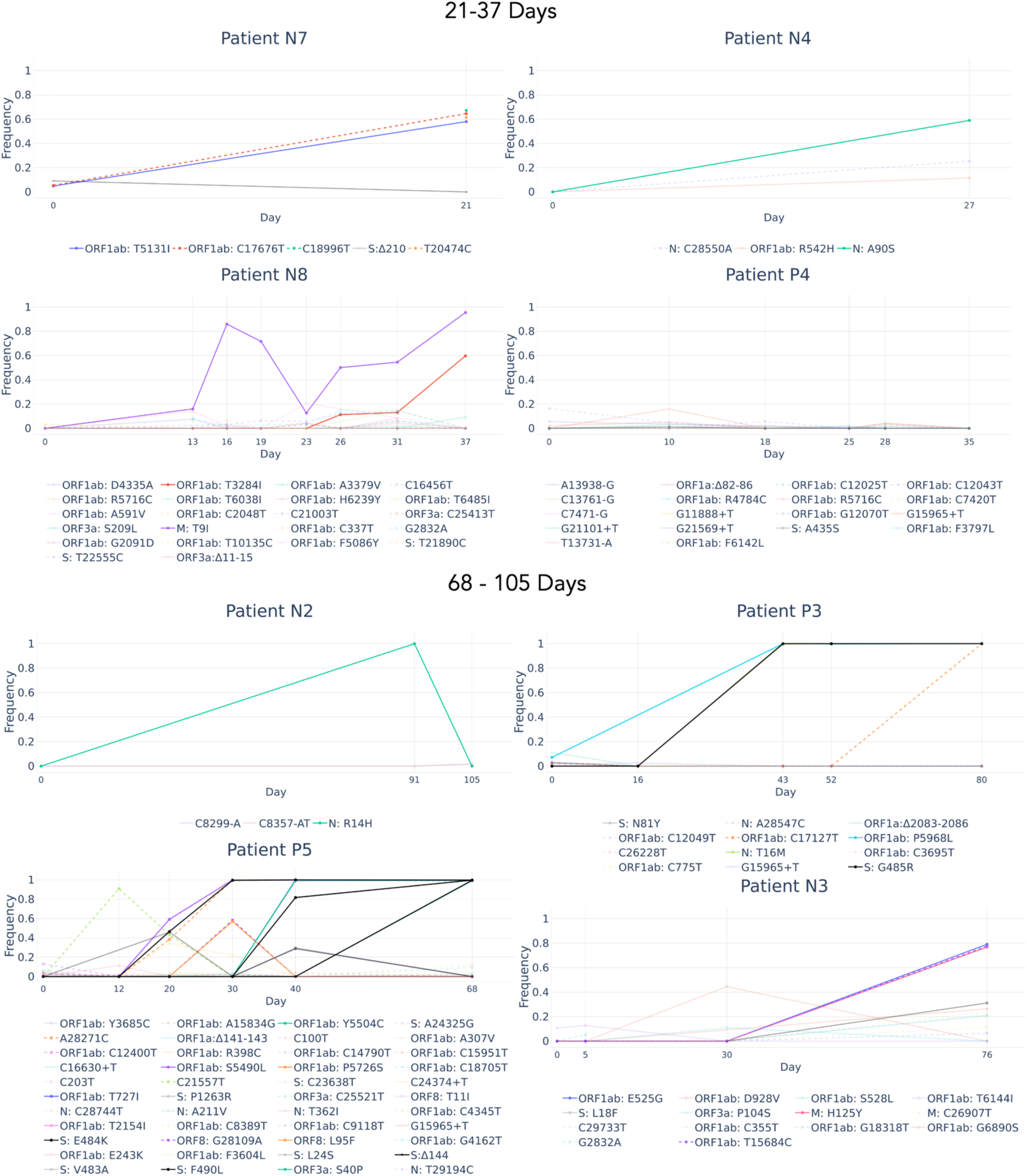
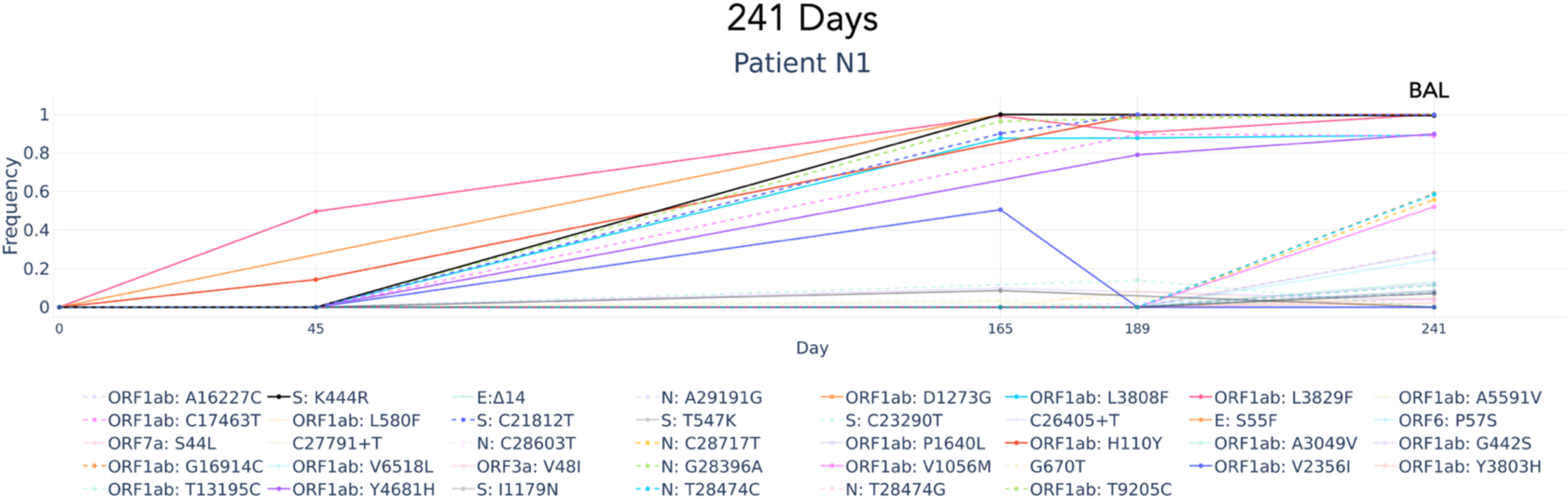
Trajectories of mutations across time. The upper panel shows four patients with a relatively short time span of sequenced infection (21-37 days), the intermediate panel show four patients with longer sequenced infections (68-105 days), and the lower panel shows one patient with a very long sequenced infection (241 days). Of note, the last sample for this patient was derived from a bronchoalveolar lavage (BAL) swab. Non-synonymous mutations in spike at high frequencies (*f*> 0.5) are shown in black, and synonymous mutations are shown with dashed lines and labeled based on their nucleotide position. Mutations occurring at low frequencies (*f* <0.5) across all time points are displayed in faded lines.

We observed dramatically different evolutionary outcomes across the different patients. These differences were evident regardless of length of sequenced infections. Thus, for example, we observed no mutations exceeding a frequency of 20% in P4 over 35 days; yet in N4, N7 and N8 we observed multiple mutations that exceed this frequency, often over a shorter time frame. When focusing on longer time scales, we again noted large differences. In N2, one mutation fixed at time point 91 and then dropped to a frequency of ∼5% two weeks later. This pattern is suggestive of the decline and rise of a subpopulation of viruses and has been noted previously (Kemp, et al. 2021; Harari, et al. 2022).

On the other hand, in P5 for example much more dramatic evolution is evident, with multiple mutations fixing sequentially. Moreover, many of these mutations are non-synonymous spike mutations, strongly suggestive that there is dramatic adaptive evolution occurring in this patient. These two opposing patterns were further exemplified in N1, the patient with the longest sequenced infection herein. For the first 45 days, very little evolution occurred. Yet by day 165, eight mutations reached near fixation, and between days 165, 189 and day 241 there were additional dramatic changes in mutation frequencies. Of note, day 241 of sequencing is derived from a bronchoalveolar lavage (BAL) sample. Interestingly, there were several mutations that rose in this sample, forming a small sub-population, as compared to time point 189; however, it is not clear if this subpopulation was unique to the BAL sample, as we do not have parallel sequencing from a nasopharynx sample at time point 241 (the latter sample was negative for SARS-CoV-2).

We next focused on the synonymous and non-synonymous (ns) divergence across time (Methods). The synonymous divergence rate, which relies on mutations that are presumable mostly neutral, is expected to yield the viral mutation rate. To obtain a more reliable estimate, we grouped all samples together (Fig. 3) and the synonymous divergence rate was estimated at 1.9×10^−6^ mutations/site/day. This value is comparable and slightly higher than an estimate of 1.3×10^−6^ ± 0.2 ×10^−6^ mutations/site/day obtained from tissue culture experiments (Amicone, et al. 2022). This value also held (albeit slightly decreased) when limiting to samples < 100 days and < 50 days (1.5×10^−6^ and 1.7×10^−6^, respectively). We further removed P5 samples where we later show there is dramatic positive selection and possible hitchhiking, and the synonymous divergence rate remained at 1.8×10^−6^.

**Figure 3.**
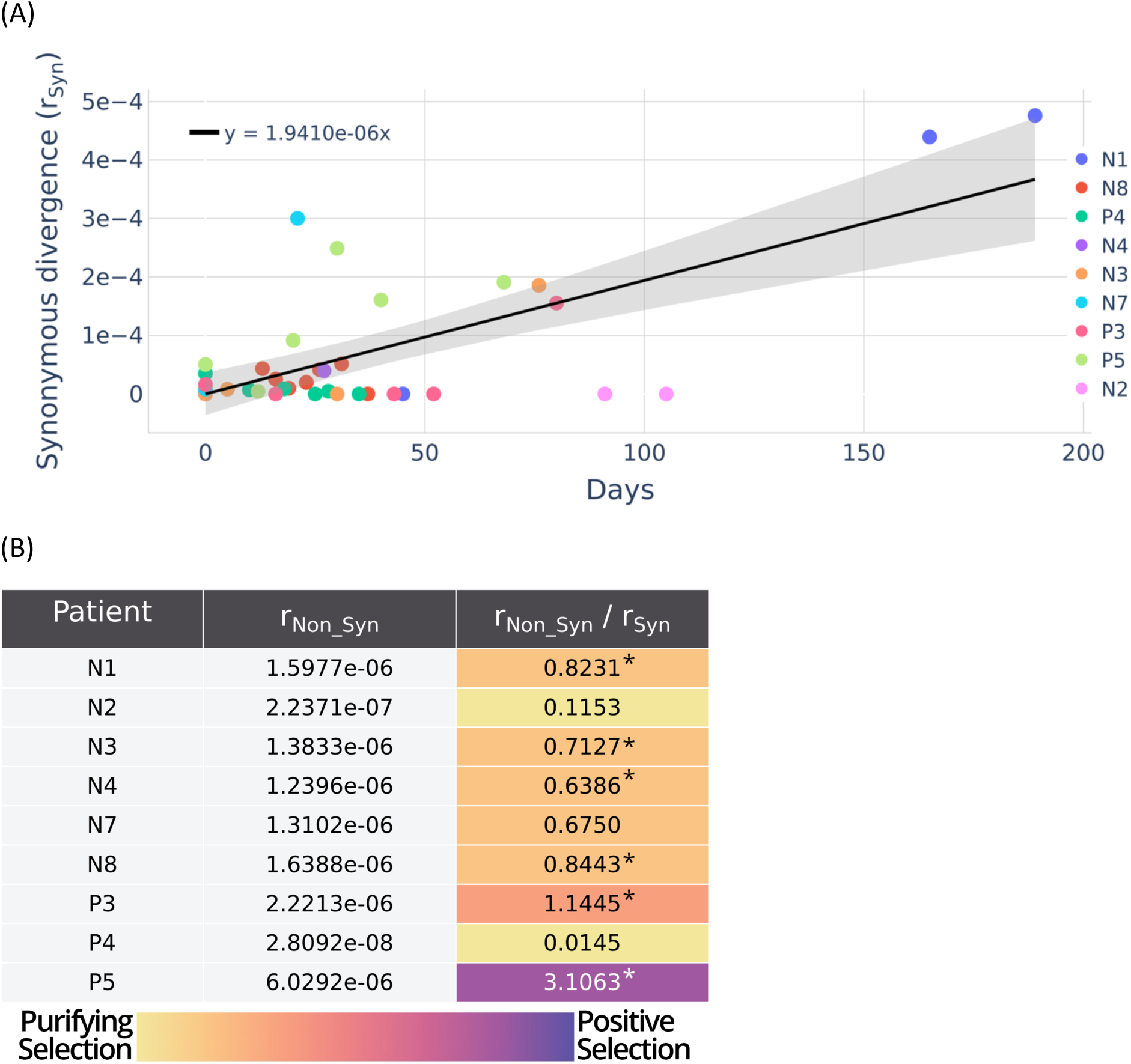
Divergence across time. (A) Synonymous divergence group across all patients, (B) Non-synonymous divergence per patient. The third column represents the ratio between non-synonymous divergence and synonymous divergence, with colors indicating the type of selection: purple for dramatic positive selection and yellow for purifying selection. Numbers marked with asterisks indicate statistically significant fits, with p< 0.01.

When focusing on the ns divergence rates, in line with the mutation trajectories shown above, we observed large variation in these rates across the samples. We noted non-significant fits for patients, P4, N2, and N7. The latter two have only two time-points. P4 has an ns rate that is effectively zero. N2 has a presumable sub-population at time point 91, but has an effective rate of zero between time 0 and time 104. Five other patients display ns rates that are roughly equal to or lower than the inferred mutation rate, and one patient, P5, stands out with a ns rate of 6×10^−6^ mutations/site/day. Thus, the only patient where we see evidence for dramatic evolution is P5, consistent with three non-synonymous spike mutations that fix in this patient. In some other patients, the ns rate is similar to the synonymous rate (e.g., P3, N1, N8), which may suggest positive selection, but this cannot be classified as dramatic adaptive evolution. We go on to explore the two opposing patterns we obtain in P4 versus P5.

In P4, we did not observe any mutations that exceeded 20%. However, when focusing on mutations at frequencies lower than 20%, we noted eighteen mutations that fluctuated at low frequencies, mostly around 1-5%. Notably, nine of these eighteen mutations were insertions or deletions (indels), and eight were point indels at coding regions, which means they disrupt the reading frame. As they occur at different genes and are quite remote, we rule out the notion that some of the indels may restore a disrupted reading frame. We thus suggest that these indels represent defective viruses present in this patient. Defective viruses are expected to segregate at frequencies similar to the mutation rate (due to mutation selection balance), and thus it is surprising to see the indels at such frequencies. One possibility is that these defective viruses thrive due to co-infections, suggesting a high viral load and high multiplicity of infection (MOI). We did not note outstanding low Ct values in this patient (Table S1).

We next move on to focus on P5, where there is a very high level of ns divergence. We have previously described this patient (Harari, et al. 2022), infected with a non-VOC, in which three ns spike antibody-evasion mutations (S:E484K, S:F490L, S:Δ144) fixed consecutively at days 30,40, and 68 (respectively). Notably the first mutation S:E484K fixed before any treatment was given to this patient, whereas S:F490L and S:Δ144 fixed after convalescent plasma treatment given at days 33 and 40-43. Overall, this suggests that the partially functioning immune system of the patient, sometimes coupled with treatment, drove adaptive evolution in this patient. Interestingly, we noted that at time point 20, two immune-evasive mutations co-existed: S:V483A (f=0.46) and S:E484K (f=0.47). They did not occur on the same reads. By time-point 30, S:V483A disappeared and S:E484K fixed, suggesting clonal competition between the two with S:E484K emerging as the winner.

## Discussion

Here, we have characterized the intra-host genetic diversity of a cohort of nine SARS-CoV-2 chronically infected individuals. Using duplicate sequencing allowed us to obtain a detailed examination of mutational dynamics during the intra-host evolution of SARS-CoV-2 during relatively long-term infections. This has allowed us to obtain several insights, elaborated on below.

First, we estimated the synonymous divergence rate of the virus during chronic infections at 1.9×10^−6^ mutation/site/day. This value is slightly higher than values obtained in tissue culture (1.3 ×10^−6^ ± 0.2 × 10^−6^) (Amicone, et al. 2022), and very similar to the rate of substitution inferred from acute infections (7.5×10^−4^ substitutions per year, which are 2×10^−6^ substitutions per day) (Neher 2022). When observing the synonymous divergence values obtained (Fig. 3A), we noted that there are many zero or near zero values that characterize the graph at early time-points and reside below the regression line. There are three likely explanations for this. First, many mutations may segregate below the 1% threshold which we use for calling. Second, given technical challenges associated with clinical samples, combined with our stringency in requiring high coverage and replicate agreement, many low frequency mutations may be overlooked. Third, from a biological perspective, genetic drift may prevail on low copy numbers of mutations, which occurs during early stages of the birth of new mutations. Conversely, we also noted many divergence values reside above the regression line, and this may be due to hitchhiking of neutral mutations that occurs as an adaptive mutation spreads. In fact, for P5 this seems to be the case, as we see synonymous mutations that rose concurrently with known adaptive spike mutations. This overall suggests that our analysis of the mutation rate is affected by two opposing forces that do not necessarily cancel each other out, and this may lead to inaccuracy in exactly estimating the mutation rate. Notably, these problems hold for previous estimates of the mutation rate as well (Amicone, et al. 2022; Neher 2022).

We go on to discuss our results on varying rates of non-synonymous divergence observed across the cohort here. We first note that our cohort was small, consisting of only nine patients. However, two recent papers have found similar results (Ghafari, Kemp, et al. 2024; Smith, et al. 2024), albeit when focusing on non-synonymous substitutions rather than on low-frequency mutations as performed herein. Notably, the work of (Smith, et al. 2024) highlighted the importance of accounting of viral sub-populations, something we do not account for here. Reassuringly, Fig. 2 suggests that such sub-populations were only observed in N2 and possibly in P5 (mutation frequencies that appear as spikes).

Notably, contrasting the non-synonymous to synonymous rate, as reported herein, is a widely used measure to detect diversifying positive selection, but it will usually capture only dramatic adaptive evolution, and may fail to pick up episodic positive selection. Thus, the fixation of S:K444R in N1, a mutation associated with immune evasion, is likely due to positive selection. Yes, it is an isolated event over 241 days of sequenced infection, suggesting this patient did not experience dramatic adaptive evolution that likely characterized VOCs that emerged in chronic infections.

What would explain the sparsity of infections with dramatic adaptive evolution? First, there are large differences in the external selection pressures applied to each patient, i.e., the therapeutic agents each patient received, which would potentially lead to different selective pressures operating on the virus (Harari, et al. 2022). Second, we surmise that the immune state of each patient may be different. Thus, patients with the most severe immunosuppression may exert the least selective pressure on the virus, at least from the point of view of the arm of immunity that is suppressed. On the other hand, patients with a weakly functioning immune system would lead to the strongest form of pressure to evolve. Overall, this would lead to large differences among patients as we and others have observed. However, to generate a transmissible variant it only takes one such patient. We note that P5 is an example of a patient that fixed four mutations at positions with lineage-defining mutations of VOCs that emerged later on (S:144, S:484, S:490, 28271). Indeed, we observe a 3X rate of non-synonymous evolution in this patient. Overall, our results thus highlight that dramatic adaptive evolution in chronic infections may be rare, yet it would only take one such rare event to spawn a new highly transmissible variant.

## Methods

### Virus genome sequencing

Virus genome sequencing was performed on residual RNA extracted from nasopharyngeal swabs and one Bronchoalveolar lavage (BAL) sample (time point 241 of N1), collected from nine chronically infected individuals who were sampled at multiple time points per patient (Table 1). All patients were immunocompromised, and some of them have been previously described (Table 1). Comprehensive clinical data was gathered and summarized in Tables 1 and S1.

All samples underwent whole-genome sequencing of SARS-CoV-2 using the ARTIC protocol (https://artic.network/ncov-2019), with a few samples undergoing minor changes in the library preparation protocol, using a Nextera DNA Flex Library Prep kit, instead of NEBNEXT Ultra II DNA Library Prep Kit. For those samples, the primer trimming technique was adjusted, as explained later on. For samples from 2020-2021 (pre-VOC or Alpha) we used the V3 primers and for samples from 2022 (denoted as BA.1 or BA.2) we used the V4.1 primers. Sequencing was performed using Illumina Miseq 250-cycle V2 kits either at the Technion Genomic Center or at the Tel Aviv Genomics Research Unit in Israel.

### Determining genome consensus sequences and lineage

Depending on the library preparation kit, raw reads were trimmed from the various primers that were used in the multiplex PCR using pTrimmer (Zhang, et al. 2019) or manual trimming. Mapping and variant calling was performed using either our in-house pipeline AccuNGS (Gelbart, et al. 2020), or using iVar (Grubaugh, et al. 2019). Results were very similar and were used interchangeably, although since we noted iVar was a little more stringent, it was used for most analyses unless otherwise noted. Since AccuNGS reports more data, it was useful for detection of unreliable mutation inferences.

For each patient, the consensus sequence of each first time-point was determined based on sites with coverage of at least 10×. Only mutations with a frequency *f* of at least 80% were introduced into the consensus sequence. Loci with mixed populations at frequencies lower than 80% were considered ambiguous and marked by ‘N’. The ‘Pangolin’ network was used to identify the consensus sequence lineage of all the samples (https://cov-lineages.org/resources/pangolin.html) (Rambaut, et al. 2020), and rule-out re-infections.

### Inference of mutation frequencies given duplicate sequencing

To infer mutations at mid to low frequencies (i.e., *f* < 80%), we sequenced all samples in duplicate, from the RNA extraction process and onwards. To account for duplicates while inferring mutations, we created a simple computational pipeline described below that was applied.

For each patient, the consensus sequence of the first time-point serves as a reference for the next time-points. Our filtering pipeline consisted of two phases:

Phase 1 – processing each sample without accounting for its duplicate. In this phase, we filter for mutations with coverage depth >= 100x, base count (alternative depth) >= 50x, and *f* > 0.01. mutations that fail one of these filters are assigned an NA value.

Phase 2 – processing each sample while accounting for its duplicate. In this phase, we filter out mutations that do not have a strong agreement between frequencies *f_1_* and *f_2_* of both duplicates, as depicted in the decision tree in Fig. 4. We allow for a stronger disagreement between high frequencies (Δ=|*f_1-_ f_2_*| <= 0.3) as compared to low frequencies (Δ <= 0.1). High or low frequency mutations are those for which *f_1_* , *f_2_* are higher or lower than 0.5, respectively. A disagreement leads to a mutation assigned an NA value, whereas an agreement leads us to calculate weighted averages of the two frequencies *f_1_* , *f_2_* , with coverage used as weights, to assign a final frequency to each such mutation.

**Figure 4.**
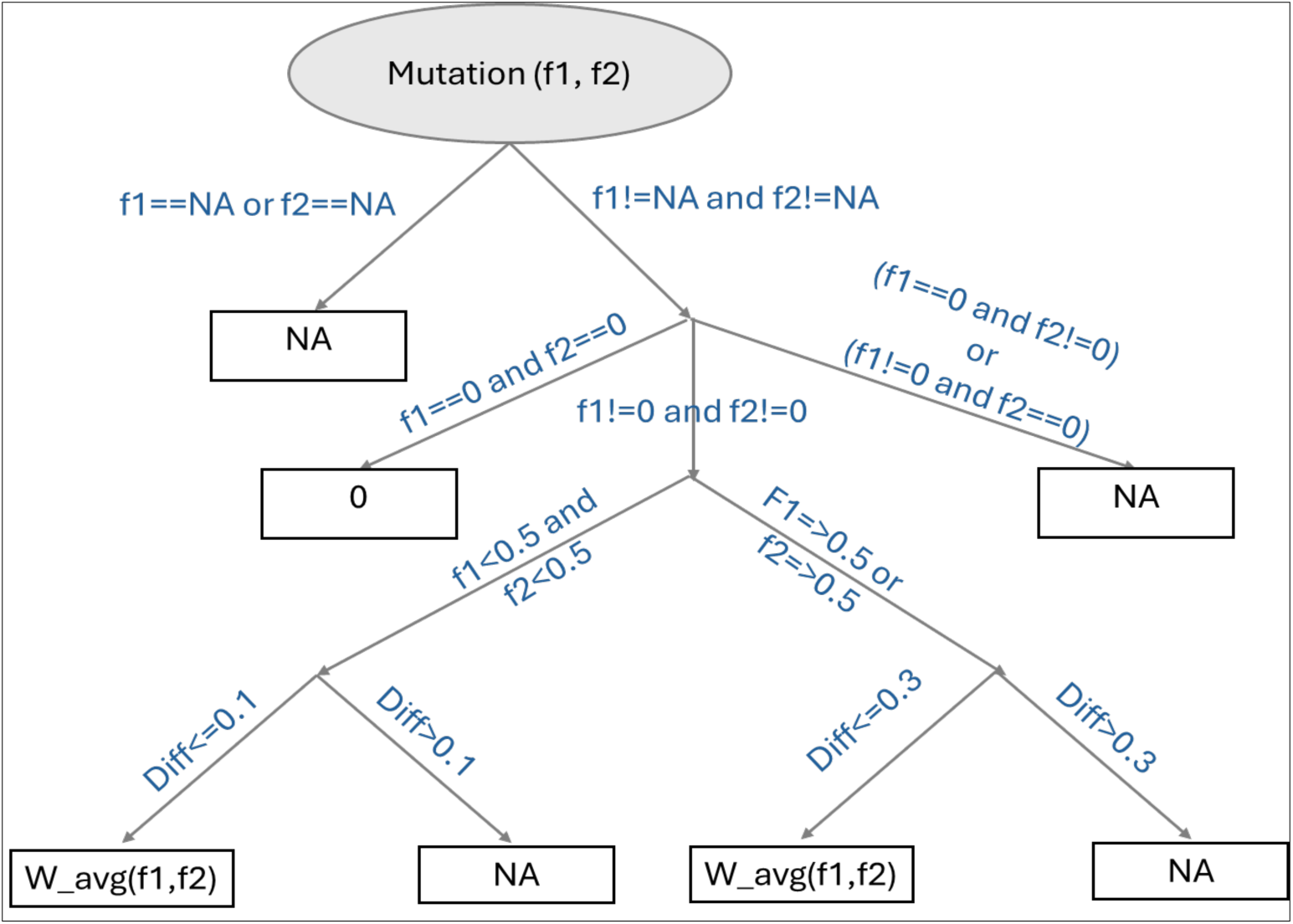
Phase 2 filtering decision tree. *f*_1_ and *f*_2_ are frequencies of a given mutation from each of the duplicates.

### Pi diversity

The nucleotide *π* diversity was calculated as described previously (Zhao and Illingworth 2019).

*π* diversity was inferred per position using the consensus sequence base as the major variant and the highest frequency mutation as the minor variant (if present, other mutations were removed, and remaining frequencies were normalized to sum of one). The final *π* diversity values per sample were determined as the average rate across all loci in the genome, using the filtering pipeline described above. Fig. 1 is based on AccuNGS results, similar results were obtained with the iVar data.

### Suspicious mutations

Our initial analysis pointed out that some mutations were recurrent across samples and seemed more like sequencing errors (Fig. S5). However, since our cohort was relatively limited in size, we set out to infer suspicious mutations using a larger cohort of acutely infected individuals previously published (Bendall, et al. 2023). The cohort consisted of various SARS-CoV-2 variants of concern (VOCs): 25 patients were infected with a non-VOC virus, 19 infected with Alpha, 2 infected with Gamma, 26 infected with Delta and 38 infected with Omicron. Samples whose VOC was unspecified were excluded from this analysis. Similar to herein, all samples had been sequenced in duplicate from the RNA extraction step onward. Alignment, consensus generation, and mutation inference were performed as described above. As we were interested in the identification of previously undetected false mutations, a mutation did not need to surpass any base count or position coverage depth minimal thresholds, nor to be present in both sequencing replicates. However, we focused our analysis on mutations with 0.01 ≤ *f* ≤ 0.2 .

Focusing on VOCs for which we had more than 10 samples, we searched for mutations that recurred across more than 20% of each VOC’s samples. For characterization purposes, we report these mutations in a detailed table (available on Zenodo), including parameters of interest for each mutation, such as: the number of samples from each VOC containing it, the CDS that it occurred in (or alternatively, if it is located within 5’UTR, 3’UTR or a TRS region), the %GC content in the 50 nucleotides surrounding it, its distance from the closest primer of each ARTIC primer set, and fitness estimations from acute infection transmission chains (Bloom and Neher 2023). We examined the single and pairwise distributions of different parameters, with the aim of understanding whether suspicious mutations are biological or rather technical errors (Figs. S4, S5).

Finally, suspicious mutations were masked from all the chronic samples herein.

### Divergence rates

Divergence values are calculated similar to (Xue and Bloom 2020) by first summing over all mutation frequencies of a certain type (non-synonymous or synonymous) at a specific time-point, including both minor and major variants, and excluding indels. As described above, mutation frequencies are always measured as compared to the consensus sequence at the first time point sequenced. To obtain a per site non-synonymous or synonymous divergence value, we divided by the number of possible non-synonymous or synonymous mutations that can be created. Based on the Wuhan reference genome, a total of 29,264 bases are part of coding regions, *n*=21070 (72%) of point mutations at coding regions are non-synonymous, and *n*= 6438 (22%) are synonymous (the rest create premature stop codons). Thus, non-synonymous or synonymous divergence values *d* are calculated respectively as:

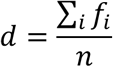

Divergence rates are calculated by performing linear regression of divergence values against time (defined for each sample as time from first time point sequenced) with the intercept set to zero. Similar results were obtained when refraining from setting the intercept to zero.

## Ethics statement

This study was approved by the Tel Aviv Sourasky Medical Center ethics board (Helsinki approval numbers 1042-20-TLV and 0021-22-TLV). It was further approved by the Tel Aviv University IRB (approval numbers 0004435-1 and 0004708-1). Requirement for informed consent was waived given the retrospective nature of the study.

## Code availability

All code used for calculations, analysis and graphing is available in the repository https://github.com/Stern-Lab/chronic-time-series-2024

## Data Availability

All sequencing data presented in this paper are available in the sequencing read archive (SRA). Previously published data (replicate 1 of P3, P4 and P5) are available under under BioProjects PRJNA803960. All other sequencing data are available under BioProject PRJNA1188568. Processed data including post-analysis mutation frequencies and haplotypes, as well as a list of all suspicious mutations, are available in the Zenodo database under accession code 10.5281/zenodo.14204007.

## Acknowledgements

We sincerely thank Daniel Weissman, Michael Martin and Katharina Koelle for valuable discussions. This study was supported by an ERC starting grant 852223 (RNAVirFitness) and an Israeli Science Foundation grant 1930/22 to A.S. This study was also supported by a fellowship to A.BZ, B.J., and S.H from the Edmond J. Safra Center for Bioinformatics at Tel Aviv University.

