## Supplementary material for "Diverse patterns of intra-host genetic diversity in chronically infected SARS-CoV-2 patients"

Supplementary material for Rutsinsky et al.

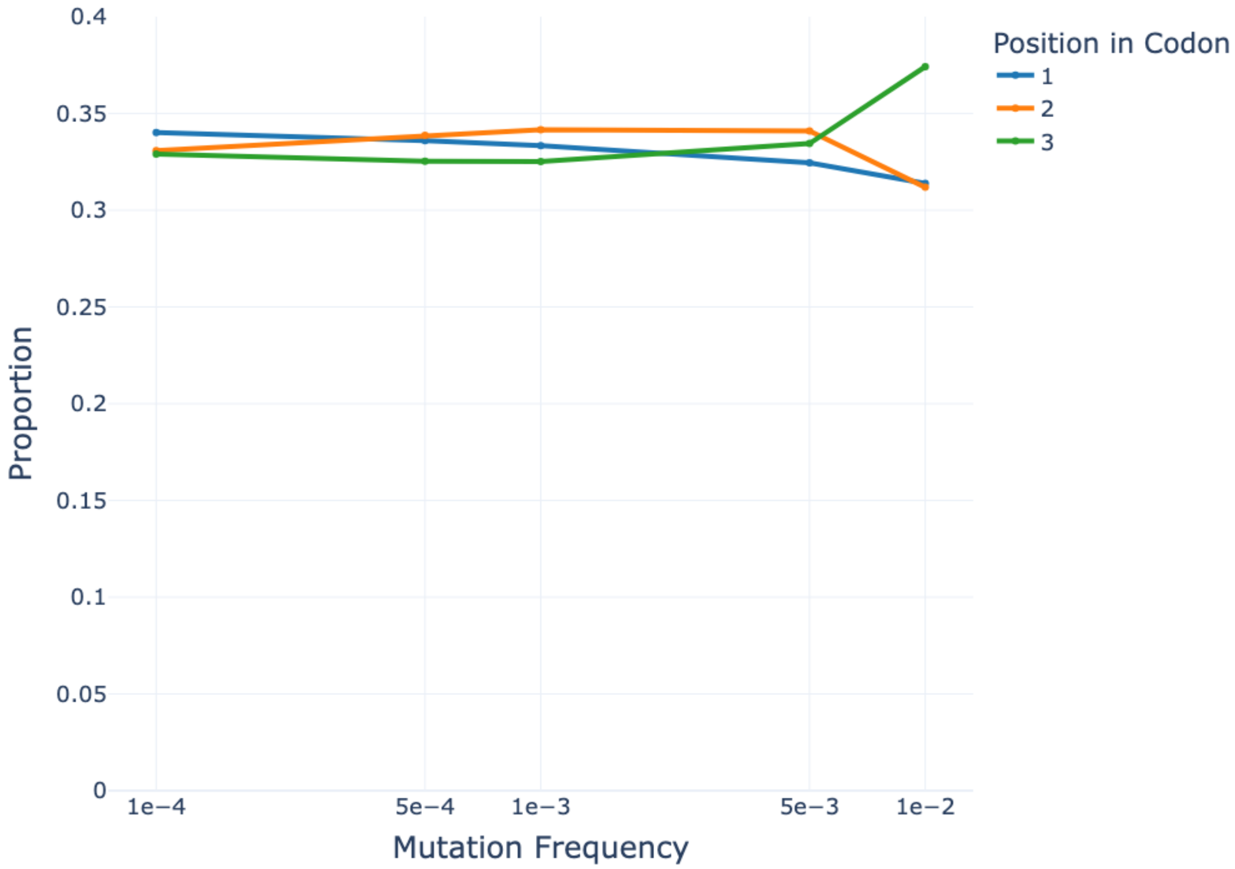
Position in Codon
1
2
3
Proportion
0.4
0.35
0.3
0.25
0.2
0.15
0.1
0.05
0
1e−4
5e−4
1e−3
5e−3
1e−2
Mutation Frequency

Figure S1.

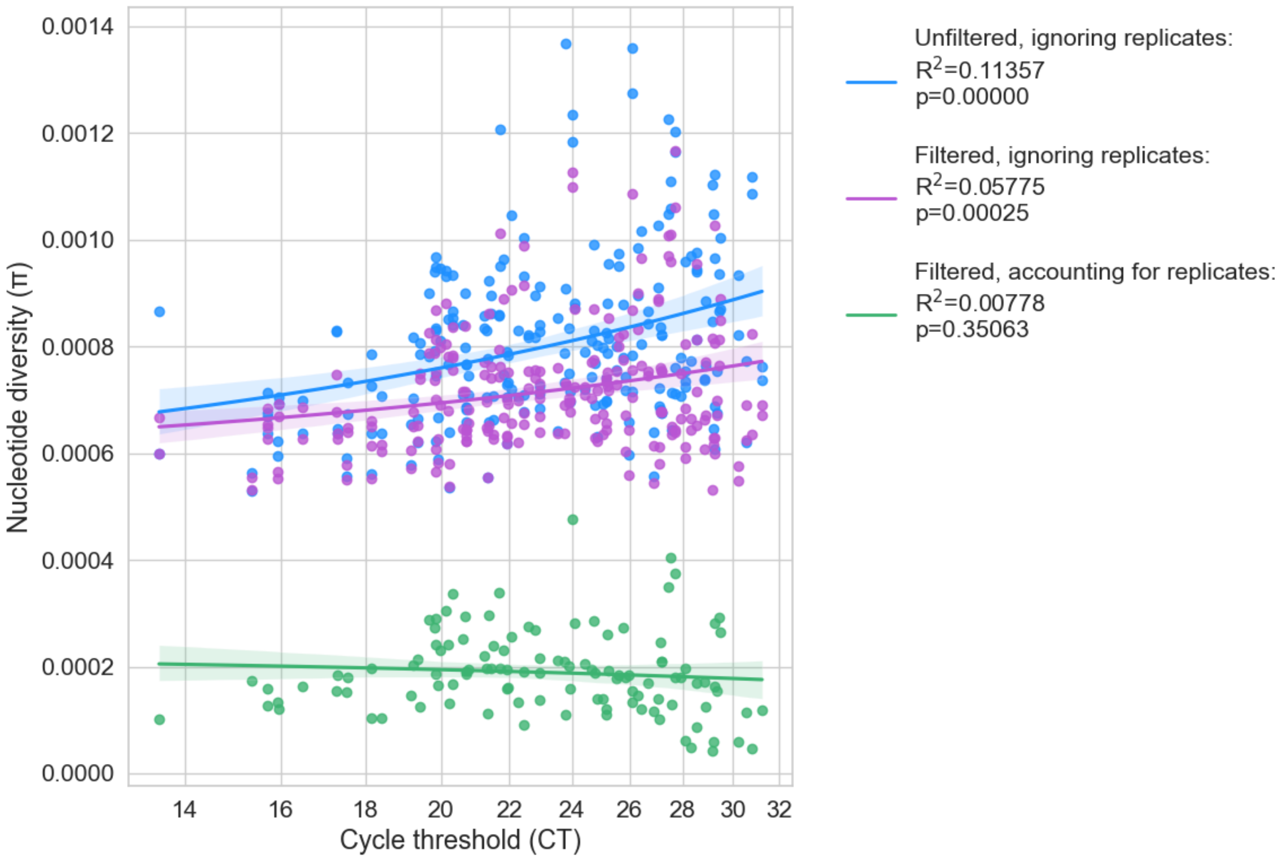


**Figure S2. Use of replicate sequencing reduces sequencing errors in acute infection samples**. Similar to Fig. 1 (shown for chronic infections), shown is the relationship between Ct, which is an inverse measure of viral load, and pi diversity of the sample in acute infections. The association between Ct and diversity disappears when mutations are called based on their presence in replicate sequencing data.

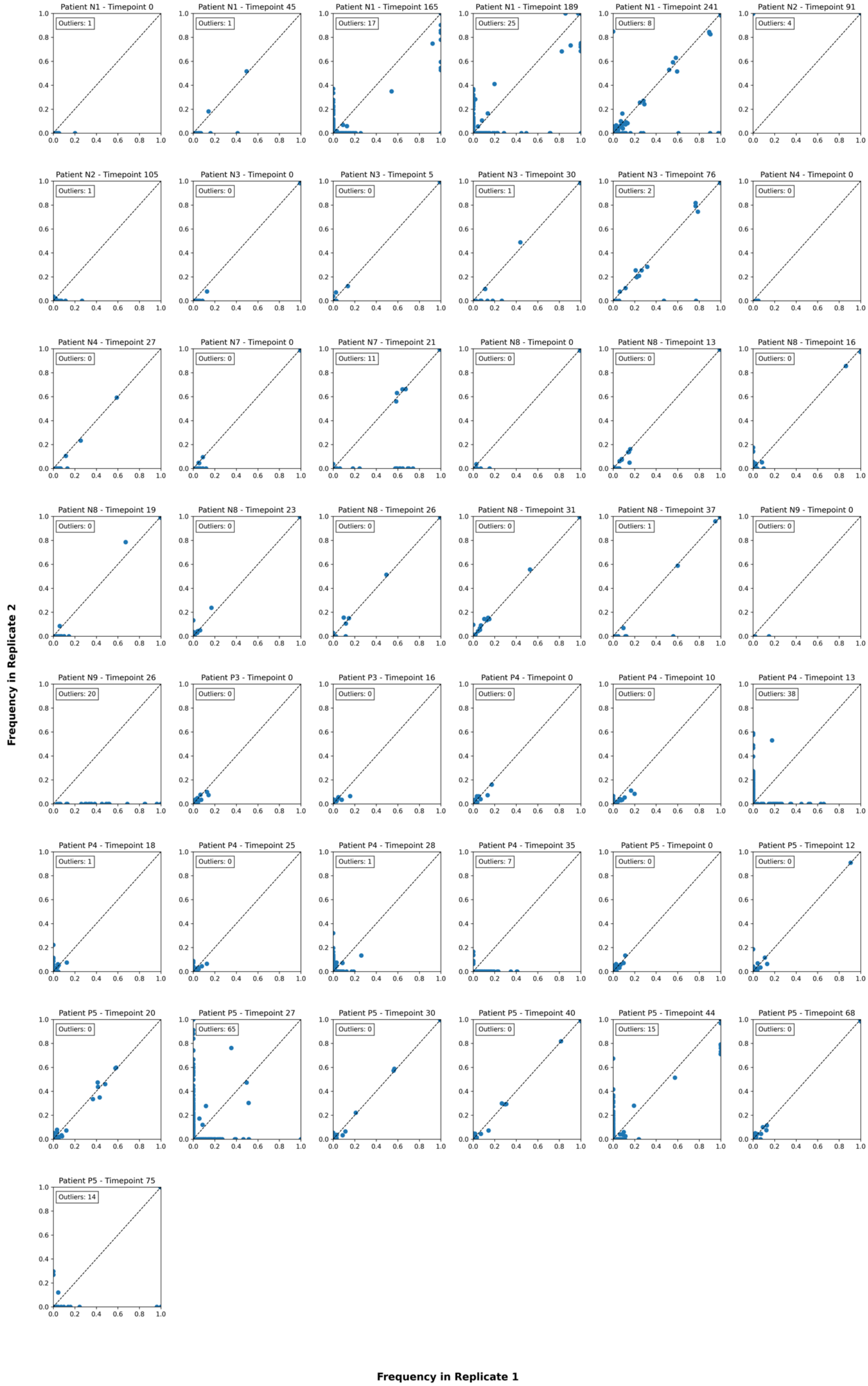
Patient N1 - Timepoint 0
Outliers: 1
Patient N1 - Timepoint 45
Outliers: 1
Patient N1 - Timepoint 165
Outliers: 17
Patient N1 - Timepoint 189
Outliers: 25
Patient N1 - Timepoint 241
Outliers: 8
Patient N2 - Timepoint 91
Outliers: 4
Patient N2 - Timepoint 105
Outliers: 1
Patient N3 - Timepoint 0
Outliers: 0
Patient N3 - Timepoint 5
Outliers: 0
Patient N3 - Timepoint 30
Outliers: 1
Patient N3 - Timepoint 76
Outliers: 2
Patient N4 - Timepoint 0
Outliers: 0
Patient N4 - Timepoint 27
Outliers: 0
Patient N7 - Timepoint 0
Outliers: 0
Patient N7 - Timepoint 21
Outliers: 11
Patient N8 - Timepoint 0
Outliers: 0
Patient N8 - Timepoint 13
Outliers: 0
Patient N8 - Timepoint 16
Outliers: 0
Patient N8 - Timepoint 19
Outliers: 0
Patient N8 - Timepoint 23
Outliers: 0
Patient N8 - Timepoint 26
Outliers: 0
Patient N8 - Timepoint 31
Outliers: 0
Patient N8 - Timepoint 37
Outliers: 1
Patient N9 - Timepoint 0
Outliers: 0
Patient N9 - Timepoint 26
Outliers: 20
Patient P3 - Timepoint 0
Outliers: 0
Patient P3 - Timepoint 16
Outliers: 0
Patient P4 - Timepoint 0
Outliers: 0
Patient P4 - Timepoint 10
Outliers: 0
Patient P4 - Timepoint 13
Outliers: 38
Patient P4 - Timepoint 18
Outliers: 1
Patient P4 - Timepoint 25
Outliers: 0
Patient P4 - Timepoint 28
Outliers: 1
Patient P4 - Timepoint 35
Outliers: 7
Patient P5 - Timepoint 0
Outliers: 0
Patient P5 - Timepoint 12
Outliers: 0
Patient P5 - Timepoint 20
Outliers: 0
Patient P5 - Timepoint 27
Outliers: 65
Patient P5 - Timepoint 30
Outliers: 0
Patient P5 - Timepoint 40
Outliers: 0
Patient P5 - Timepoint 44
Outliers: 15
Patient P5 - Timepoint 68
Outliers: 0
Patient P5 - Timepoint 75
Outliers: 14
Frequency in Replicate 2
Frequency in Replicate 1

**Figure S3**. Scatterplots of duplicate sequencing per patient sample. There was concordance in most samples, yet some samples displayed strong discordance. When samples from similar timepoints were available, such samples were removed from the analysis (P4 t=13, P5 t=27,44,75). The number of outliers represents the mutations where the frequency exceeds 0.2 in one replicate but is 0 in the other

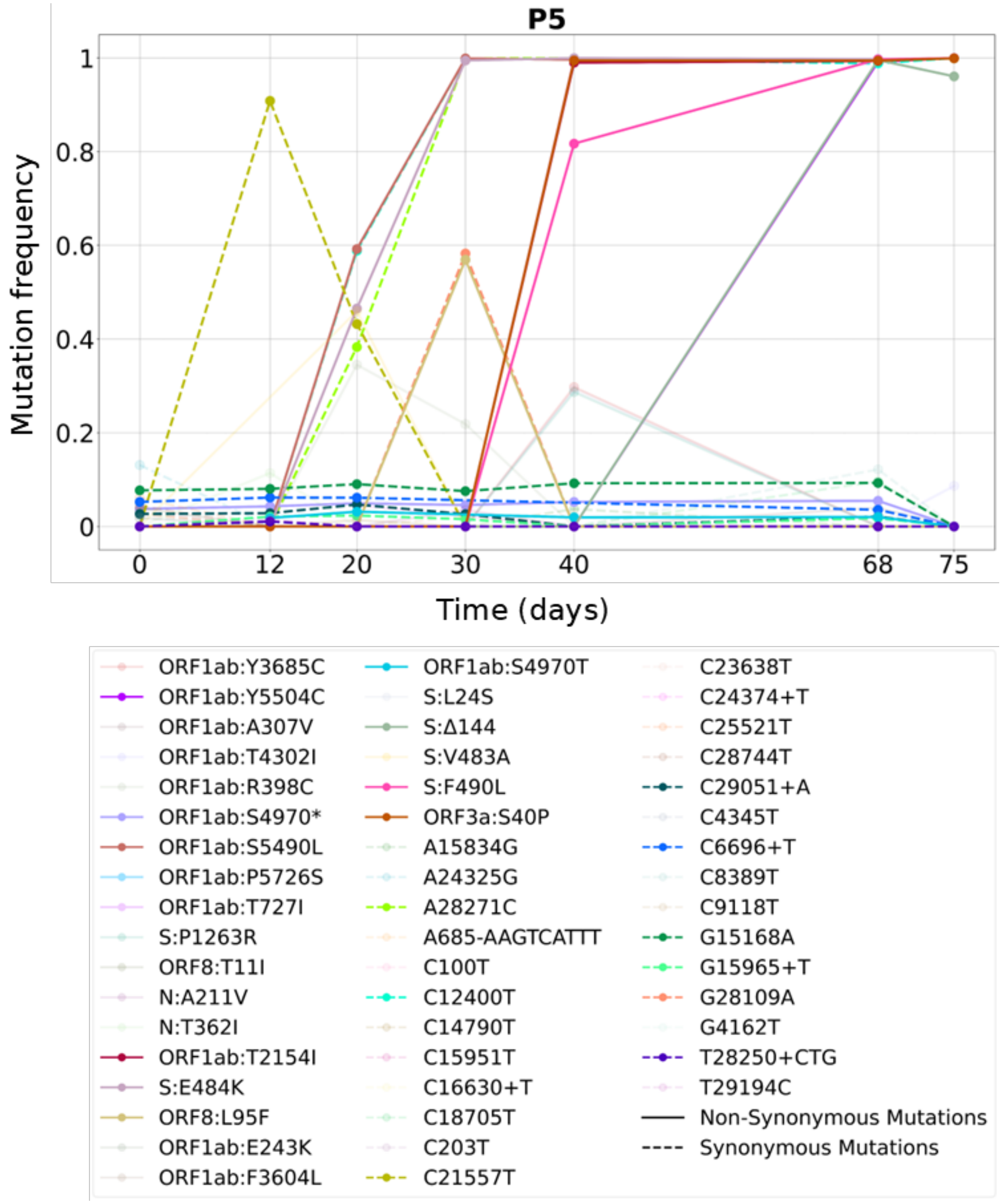


**Figure S4.** Example of mutation frequencies across time without exclusions of suspicious mutations in P5. Despite several mutations fixing across time, we note the constant presence of many mutations (e.g., C6696+T, ORF1ab:S4970T, ORF1ab:S4970*). All these mutations were subsequently found to occur repeatedly in many samples of acute infection SARS-CoV-2, and were filtered from our analysis.

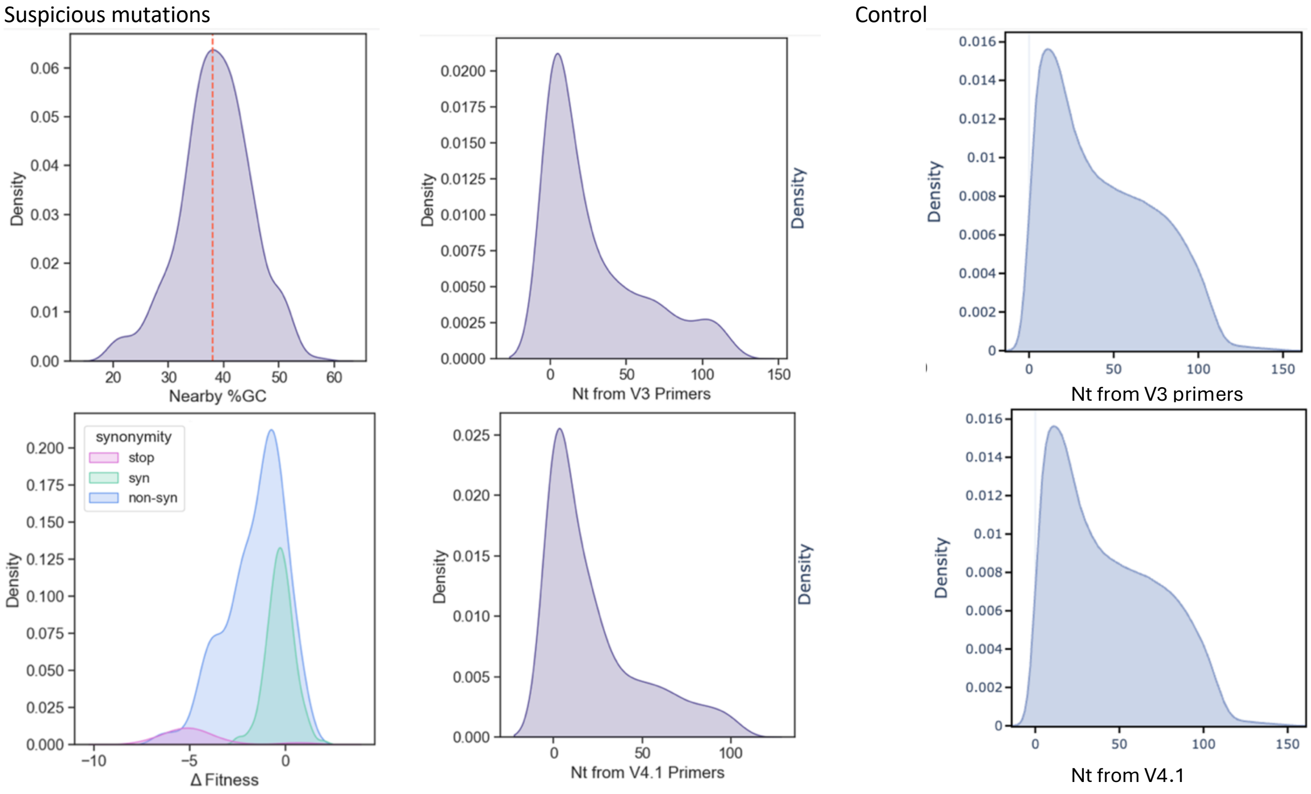


**Figure S5. Distributions of parameters examined for suspicious mutations**. (A) %GC surrounding suspicious mutations, with the dashed line representing the genome-wide average. Suspicious mutations did not tend to have lower or higher surrounding GC content. (B) Fitness of suspicious mutations (denoted as Δfitness) stratified by mutation type. No difference was observed versus genome-wide inferences of fitness (Bloom and Neher 2023). (C, D) Distances of suspicious mutations to V3 or V4.1 primers. (E, F) Genome-wide distances of all positions from V3 or V4.1 primers (control). Suspicious mutations show significantly lower distances to primers as compared to the control (Mann-Whitney, $p<10^{-7}$), suggesting many are associated with errors in priming.

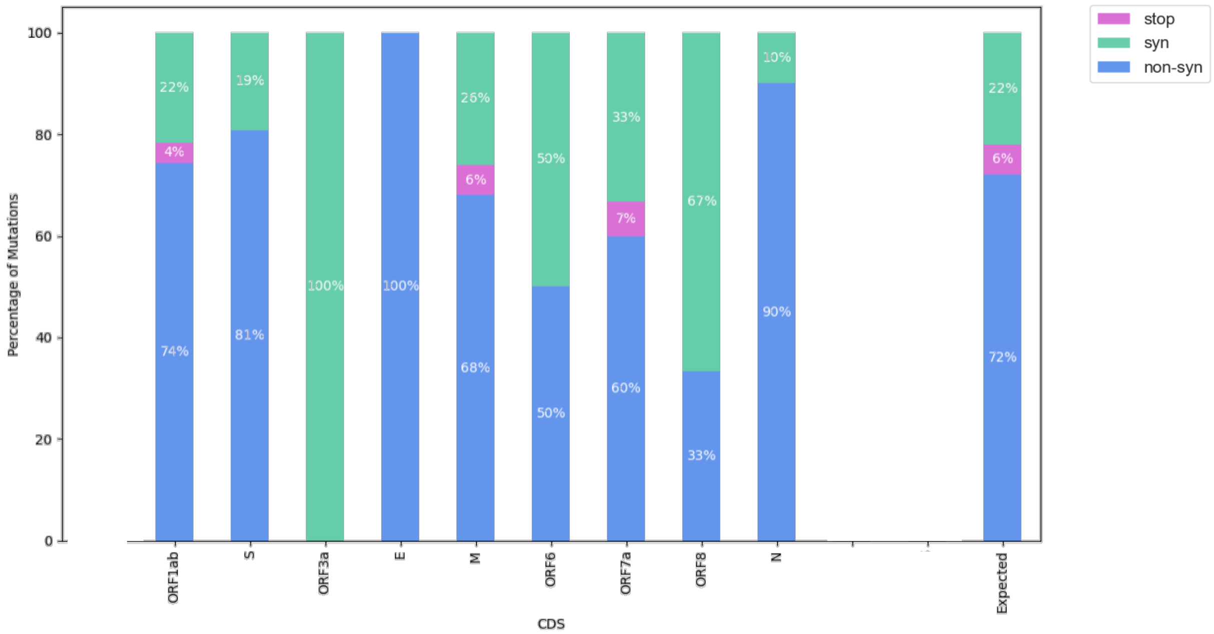


**Figure S6. Proportions of mutation effects (synonymous/non- synonymous /premature stop) of the suspicious mutations stratified by gene/genomic region.** Last column represents expected mutation effects based on single mutations on the background of the Wuhan reference genome. In the two longest open reading frames (ORF1ab and S) no significant difference was found compared to the expected counts based on a Fisher exact test.

**Table S1. Ct values for all samples of all patients**. Ct values were averaged across all marker genes used. All samples were from obtained from nasopharyngeal swabs, with the exception of N1 time point 241, which was a bronchoalveolar lavage sample.

| Sample | Patient | Time since first sample | Ct |
|---|---|---|---|
| 741140288 | N1 | 0 | 26 |
| 741149899 | N1 | 45 | 27.6 |
| 741294879 | N1 | 165 | 31.5 |
| 741296489 | N1 | 189 | 32.67 |
| 741299538 | N1 | 241 | 27.4 |
| 741210690 | N2[1] | 0 | 22 |
| 741286954 | N2 | 91 | 23.37 |
| 741288907 | N2 | 105 | 30.83 |
| 741148584 | N3 | 0 | 30 |
| 741149282 | N3 | 5 | 26.5 |
| 741284250 | N3 | 30 | 28.9 |
| 741288510 | N3 | 76 | 27.5 |
| 741149041 | N4 | 0 | 18.62 |
| 741284247 | N4 | 27 | 26.73 |
| 741146042 | N7 | 0 | 29.5 |
| 741149797 | N7 | 21 | 29.27 |
| 741146626 | N8 | 0 | 26.5 |
| 741149060 | N8 | 13 | 24.35 |
| 741149428 | N8 | 16 | 27.5 |
| 741149794 | N8 | 19 | 28.1 |
| 740925216 | N8 | 23 | 29.37 |
| 740925663 | N8 | 26 | 19 |
| 741283212 | N8 | 31 | 22 |
| 741283709 | N8 | 37 | 34 |
| 740665529 | P3 | 0 | 14.07 |
| 740656183 | P3 | 16 | 27.43 |
| 740677502 | P3[1] | 43 | 27 |
| 740812853 | P3[1] | 52 | 25 |
| 740687212 | P3[1] | 80 | 30 |
| 740652926 | P4 | 0 | 21.47 |
| 740658998 | P4 | 10 | 28.43 |
| 740663063 | P4 | 18 | 30.27 |
| 740666995 | P4 | 25 | 27.53 |

| | | | |
|---|---|---|---|
| 740668927 | P4 | 28 | 29.07 |
| 740671764 | P4 | 35 | 30.5 |
| 740605134 | P5 | 0 | 22.5 |
| 740691329 | P5 | 12 | 18.95 |
| 740696660 | P5 | 20 | 32.67 |
| 740702156 | P5 | 30 | 18 |
| 740706570 | P5 | 40 | 28.9 |
| 740694156 | P5 | 68 | 25 |
| 740628355 | P5 | 75 | 28 |

[1] Samples for which only one replica of sequencing was performed previously.
